# Planum Temporale grey matter volume asymmetries in new-born monkeys (*Papio anubis*)

**DOI:** 10.1101/2021.02.22.432274

**Authors:** Yannick Becker, Romane Phelipon, Julien Sein, Lionel Velly, Luc Renaud, Adrien Meguerditchian

## Abstract

The Planum Temporale (*PT)* is one of the key hubs of the language network in the human brain. The gross asymmetry of this perisylvian region toward the left brain was considered as the most emblematic marker of hemispheric specialization of language processes in the brain. Interestingly, this neuroanatomical signature was documented also in newborn infants and preterms, suggesting the early brain’s readiness for language acquisition. Nevertheless, this latter interpretation was questioned by a recent report in nonhuman primates of a potential similar signature in newborn baboons *Papio anubis* based on *PT* surface measures. Whether this “tip of the iceberg” *PT* asymmetry is actually reflecting asymmetry of its underlying grey matter volume remain unclear but critical to investigate potential continuities of cortical specialization with human infants. Here we report a population-level leftward asymmetry of the Planum Temporale grey matter volume in *in vivo* 34 newborn baboons *Papio anubis*, which showed intra-individual positive correlation with *PT* surface’s asymmetry measures but also a more pronounced degree of leftward asymmetry at the population-level. This finding demonstrates that *PT* leftward structural asymmetry in this Old World monkey species is a robust phenomenon in early primate development, which clearly speaks for a continuity with early human brain specialization. Results also strengthen the hypothesis that early PT asymmetry might be not a human-specific marker for the pre-wired language-ready brain in infants.

## Introduction

The majority of language processes is asymmetric in the human brain, involving a specialization of the left hemisphere (Vigneau et al., 2006). The most emblematic marker of such a language cerebral organization is the gross asymmetry of the Planum Temporale (*PT*) toward the left hemisphere. This perisylvian region, which constitutes the floor of the Sylvian fissure, posterior to Heschl’s gyrus and overlaps with Wernicke’s area, is one of the key hubs of the language network in the human brain. In fact, the left *PT* was significantly activated in a variety of language processing tasks in adults (Shapleske et al., 1999, Vigneau et al., 2006, Josse et al., 2006).

Since the first discovery that the *PT* was larger in the left hemisphere than the right in most adults (Geschwind and Levitsky, 1968), it remains unclear, whether this feature constitutes a good marker of language functional lateralization. While some studies reported no match between structural and functional asymmetry of this region (Keller, 2011; Greve, 2013), clinical studies found that atypical *PT* structural asymmetry were associated with multiple language deficits (Borovsky et al., 2007; Dronkers et al., 2004; Foundas et al., 2004; Wernicke, 1874). In addition, in a recent study, higher density of dendrites and axons in the *PT* were associated with faster neurophysiological processing of auditory speech (Ocklenburg et al., 2018). Moreover, in a second recent study, structural *PT* asymmetry was found associated with functional lateralization of an adjacent auditory area at the end of the Sylvian fissure during a language task (Tzourio-Mazoyer et al., 2018).

Interestingly, leftward PT asymmetry was detected early in the development at both the functional level in three-month-old infants in response to speech (Dehane-Lambertz et al., 2002) and at the structural level in newborn and in preterms (Witelson and Pallie, 1973; Wada, 1975; Chi et al., 1977, Dubois et al., 2010; Hill et al., 2010; Glasel et al., 2011). Such early features of language brain lateralization suggest that the infant brain might be already pre-wired for language acquisition (e.g. Dehaene-Lambertz et al., 2002).

However, the human uniqueness of structural *PT* asymmetry was questioned by studies highlighting also a population-level leftward asymmetry by *PT* surface measures in chimpanzees (Gannon et al. 1998, Hopkins et al., 1998, Spocter et al. 2020) and in baboons (Marie et al. 2018). In this latter old world monkey species, *PT* leftward surface biases were found not only in adults but also recently in newborn baboons (Becker et al., 2021, see also Xia et al., 2019 for a study in macaques using cortical surface-based morphometry), suggesting it might reflect the asymmetry of its underlying grey matter volume and is thus not specific to human early brain development. In fact, *PT* surface area measures quantified the depth of the sylvian fissure’s floor. It might be thus not excluded that the asymmetry of the sulcal surface area of this region might be an appropriate indicator of the asymmetry of the juxtaposing grey matter volume of the *PT*. This hypothesis is supported by few studies in adult chimpanzees which focused on *PT* grey matter volume asymmetry according to both ROI manual tracing (Hopkins and Nir, 2010; Lyn et al., 2011) and voxel-based morphometry (Hopkins et al., 2008), all showing consistent leftward asymmetry with *PT* surface measures.

In the present study, we further explore this hypothesis in 34 newborn baboons by quantifying the grey matter volume of the left and right *PT* from *in vivo* MRI brain scans (Becker et al., 2021). The aim of the follow-up study is thus to investigate early individual and population-level asymmetries of the *PT* grey matter volume in newborn nonhuman primates and their potential consistencies with *PT* surface asymmetries measures within the same cohort of subjects used in Becker et al.’s study (2021).

## Materials and Methods

### Subjects

Subjects ranged from 4 to 165 days of age (Mean: 32.63; SD: 6.13) and included 21 males and 14 females. (see table in supplementary methods with subjects’ details)

All monkeys are housed in social groups at the Station de Primatologie CNRS (UPS 846, Rousset, France) and have free access to outdoor areas connected to indoor areas. All subjects are born in captivity from 1 (F1) or 2 generations (F2). Wooden and metallic structures enrich the enclosures. Feeding times are held four times a day with seeds, monkey pellets and fresh fruits and vegetables. Water is available ad libitum.

### MRI Image acquisition

Structural magnetic resonance images (MRI) were collected from a sample of 35 baboons (September 2017 to March 2020). Animals were minimally anesthetized by a veterinarian; and vital functions were monitored during the scans. High-resolution structural T1-weighted brain images were obtained with MPRAGE sequences (0.4 mm isotropic, TR = 2500ms, TE = 3.01ms) with the subject in the supine position on a Siemens 3T Magnetom Prisma scanner and using two 11cm receive-only loop coils (for more detailed procedure: Becker et al., 2021). At the end of the MRI session, when fully awaked from anaesthesia, baboons were carefully put back with their mother and then transported for immediate (or delayed) reintroduction into their social groups under staff monitoring.

### Preprocessing of Anatomical MRI

Anatomical T1w images were noise corrected with the spatially adaptive nonlocal means denoising filter (Manjón et al., 2010) implemented in Cat12 toolbox (http://www.neuro.uni-jena.de/cat/) included in SPM12 (http://www.fil.ion.ucl.ac.uk/), which runs on MATLAB (R2014a). Next, each image was manually oriented using ITK-Snap 3.6 according anterior and posterior commissures plane and the interhemispheric fissure plane.

### Manual Delineation of the Planum Temporale’s grey matter volume

Manual delineation was conducted with “ANALYZE 11.0 (AnalyzeDirect)” software and following the delimitation instructions established in previous *PT* studies in nonhuman primates using MRI (e.g., Hopkins and Nir, 2010; Lyn et al., 2011; Meguerditchian et al., 2012; Marie et al., 2018; Becker et al., 2021). The delineation of the posterior edge of the *PT* is defined by the most caudal section showing the Sylvian fissure. In humans, the anterior edge of the *PT* is delimited by the Heschl gyrus, however, in baboons the Heschl gyrus is not clearly detectable, therefore to delineate the anterior edge of the *PT* here, the most anterior cut including the Sylvius Fissure was used when the insula closes completely (when the insula fissure disappears completely posteriorly). For each slice, manual tracing was conducted from the medial most point of the Sylvius Fissure, to the most lateral point, following the most ventral edge of the fissure. Next, the raters followed the grey matter to its most inferior edge of the grey/white matter boundary. When ambiguous, the imaginary prolongation of the Sylvian Fissure was used to differentiate between the grey matter of interest and the more dorsal gyrus. This step is repeated on the next cut, advancing posteriorly, until the Sylvius Fissure disappears. If the fissure forked in an ascending or descending direction, it was preferable to follow the descending one. This manual tracing was done on the coronal plane and not sagittal, as it gives the best assessment of the total depth of the Sylvius pit, which is the “ground” of the *PT*. The manual delimitation was carried out via a graphic tablet (WACOM cintiq 13HD). Out of the 256 slices included in the MRI images, the *PT* appeared in about 20 slices (Supplementary Figure 1)

For each subject, an Asymmetry Quotient (AQ) of the left (L) and the right (R) grey matter volume was computed AQ = (R – L) / [(R + L) × 0.5] with the sign indicating the direction of asymmetry (negative: left side, positive: right side) and the value, the strength of asymmetry. Further, as reported by Hopkins and Nir (2010) for humans and great apes, the AQ was also used to classify the subjects as left-hemispheric biased (AQ ≤ –0.025), right biased (AQ ≥ 0.025), or non-significantly biased “ambi” (–0.025 < AQ < 0.025).

To reduce potential observer-dependent manual tracing biases, all the *PTs* grey matter volume were traced by a rater different from the one who traced the *PT* surface in Becker et al. (2021). The rater of the present study was blind to the *PT* surface’s tracing, data and results of Becker et al. (2021). Statistics were conducted with R 3.6.1 (R Core Team (2017). R: A language and environment for statistical computing. R Foundation for Statistical Computing, Vienna, Austria. URL https://www.R-project.org/.)

## Results

### PT Grey matter volume measures

We found a significant leftward asymmetry of the *PT* grey matter volume at a group-level in 34 newborn baboons according to a one sample t-test in the 34 subjects AQ scores (see Figure 2), *Mean* AQ = −0.121, 0.169 SD; *t*(33) = −4.2, *p* < 0.0001. Categorization of individual AQ showed also a majority of leftward *PT*-biased individuals: 24 baboons exhibited a leftward hemispheric *PT* bias (70.6%) whereas 7 exhibited a rightward *PT* bias (19.6%) and 3 no *PT* bias (8.8%). The number of leftward *PT*-biased individuals was significantly greater than the number of rightward *PT*-biased according to chi-square test (χ2 = 21.94, *p* < 0.0001).

**Figure 1.**
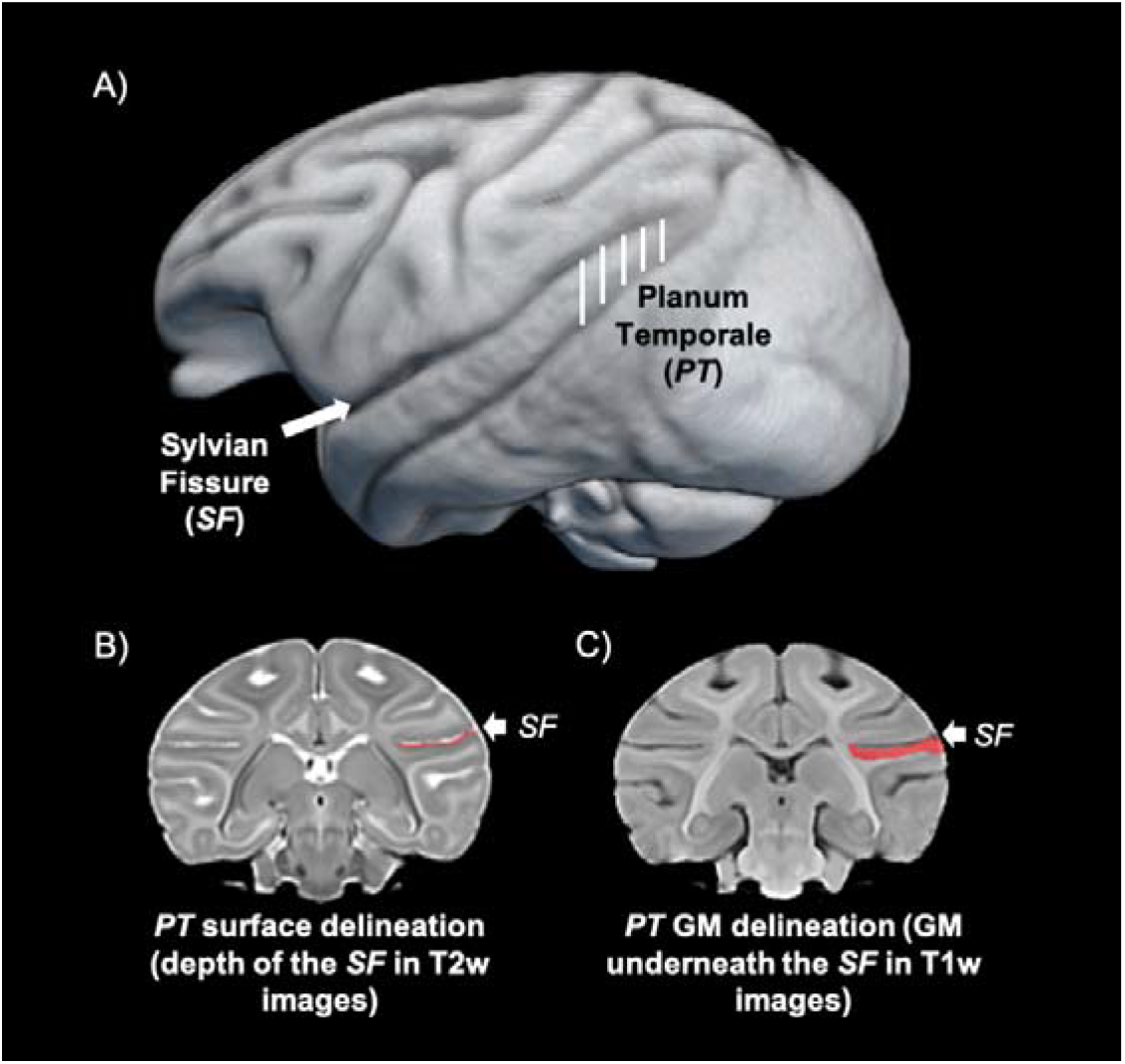
(A) 3D reconstruction of a newborn baboon brain with the *PT* region highlighted by the white lines (B) Coronal slice of the same subject with the delineation of the Sylvian Fissure’s floor (in red), used for *PT* surface measures (C)) Coronal slice of the same subject with the delineation of the grey matter (in red) underneath the Sylvian Fissure, used for *PT* volume measures.

**Figure 2.**
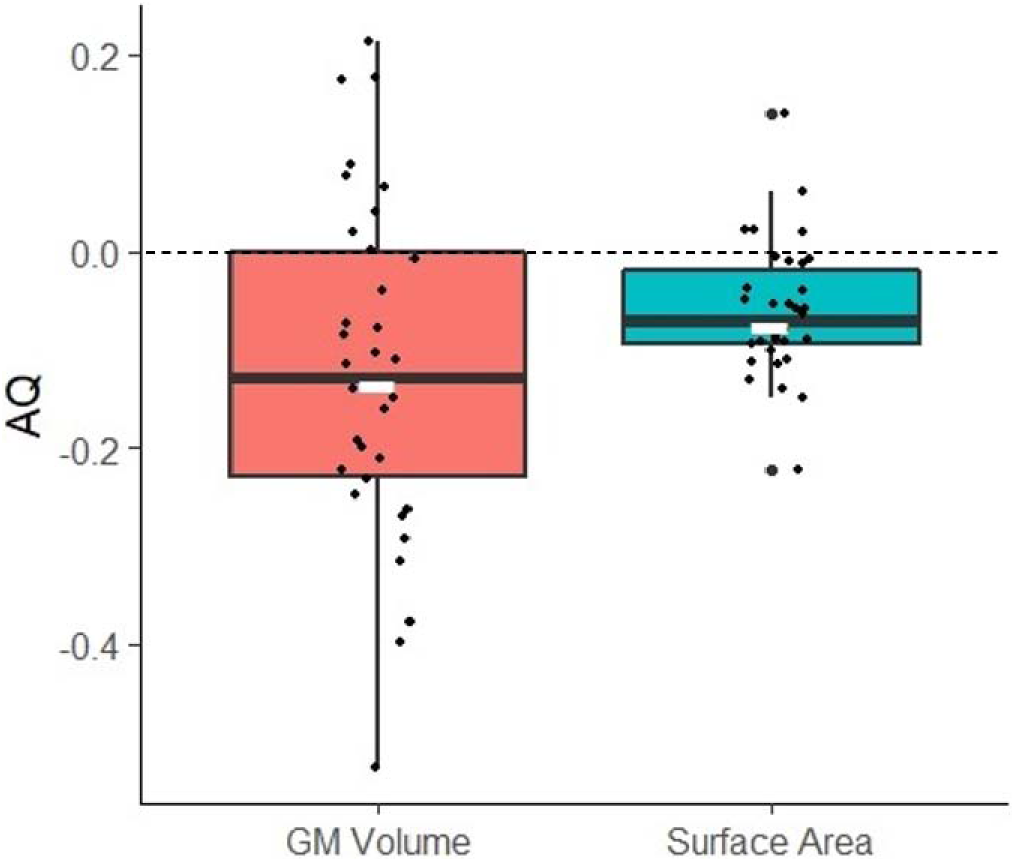
Distribution of Asymmetry Quotients (AQ) for Grey Matter (GM) volume measures in red and Surface area measures in blue for the same subjects. AQ values inferior to 0 indicate leftward lateralization, AQ values superior to 0 indicate rightward lateralization. Note the leftward lateralization for both measure types. Note also larger distribution, ie. higher AQ values for the grey matter volume measures.

**Figure 3.**
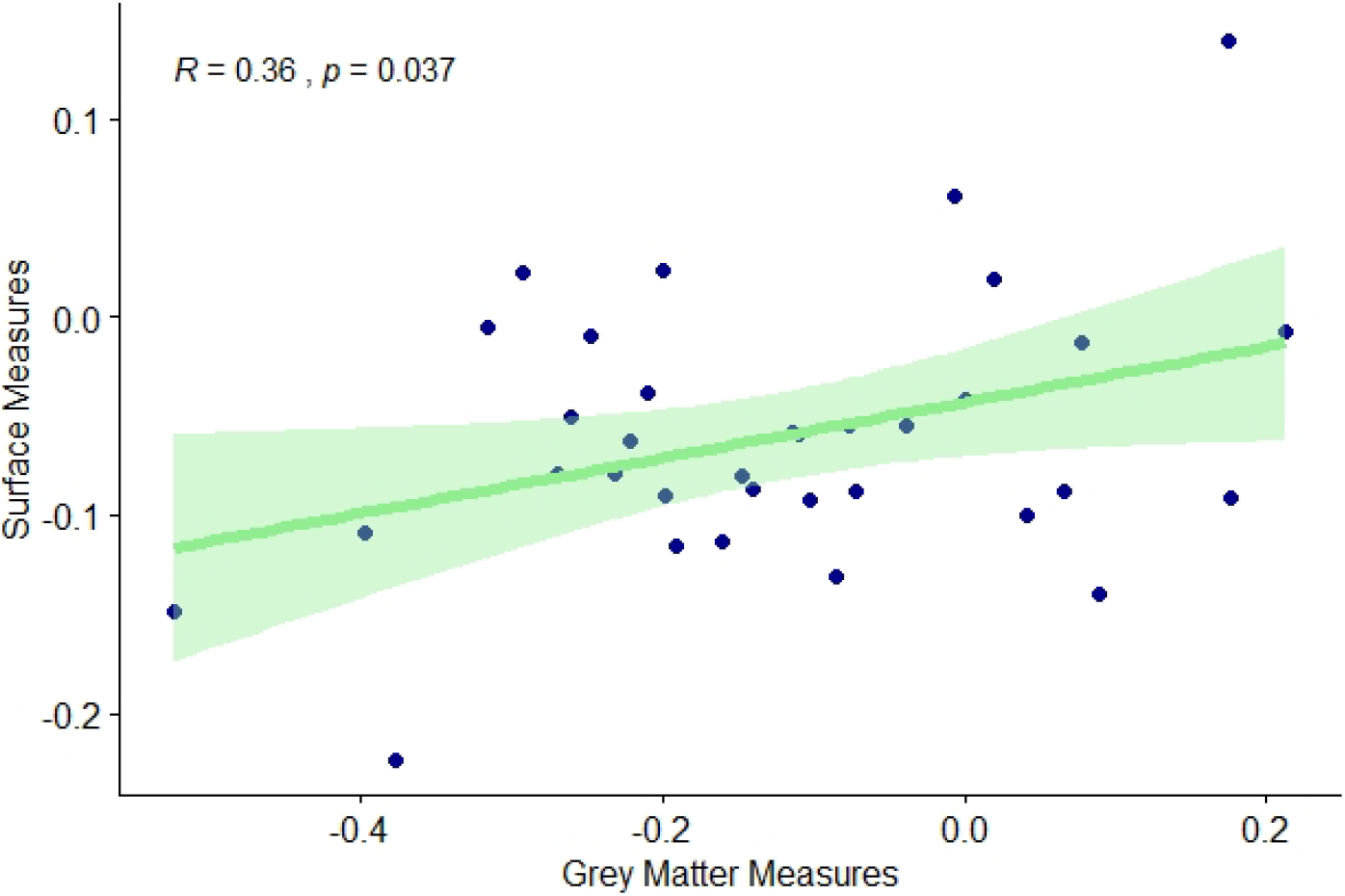
Pearson correlation between PT Surface and Grey matter volume measures.

Multiple linear regression analyses showed that the right *PT* volume (*p* < 0.001) and the left *PT* volume (*p* < 0.001) predict *PT* asymmetry strength, but not the subject’s sex, age nor brain volume.

### Correspondence between PT Surface and PT Grey Matter measures

Within the 34 individuals for whom data of *PT* surface and *PT* grey matter volume measures were independently traced by two different raters blind to the results of each other, a significant positive intra-individual correlation of AQ scores was found between *PT* surface and *PT* grey matter volume. *r*(34) = 0.36, *p* < 0.037.

In comparison to previous surface *PT* measures (Becker et al., 2021), 22 were consistent in hemispheric lateralization classification (i.e., 20 leftward, 1 rightward and 1 ambi) and 4 subjects switched direction of hemispheric *PT* bias (i.e., from leftward bias for *PT* surface to rightward bias for *PT* grey matter volume). Among the remaining 8 subjects, 6 which previously showed no significant bias (i.e. “ambi”) for the *PT* surface were found significantly lateralized for the *PT* grey matter volume and 2 which were previously classified as significantly lateralized for the *PT* surface were classified as “ambi” for the *PT* grey matter volume.

## Discussion

We find for the first time both individual and leftward population-level grey matter volume asymmetries of the Planum Temporale not only in Old World monkeys but also in a newborn non-human primate. These results showed intra-individual positive correlation with previous published *PT* surface measures on the same subjects as well as consistent leftward *PT* asymmetry (Becker et al., 2021) and suggesting *PT* surface measures may therefore reflect its underlying grey matter volume.

The distributions of individual *PT* hemispheric preferences (left, right or ambi) are quite similar between volumetric grey matter and surfacic measures, especially for the left lateralized subjects, although some inconsistency was noticed at the individual level in a minority of subjects. It remains unclear whether those variations are due to interrater-dependent variability in the measures, which leads few subjects to switch categories or to the possibility that *PT* surface measures are not entirely perfect “tip of the iceberg” predictors of the *PT* grey matter volume, especially for the subjects initially classified as ambiguously biased for *PT* surface. In fact, almost all of those latter “ambi” newborns (6 out of 7) were found to be significantly lateralized for *PT* grey volume. In addition, AQ values were overall higher in grey matter measures (AQ −0.121 12.1%) compared to surface measures (AQ −0.073 7.3%). A similar effect was found in Hopkins and Nir’s paper (2010), which showed a 4.96% larger left hemisphere when measuring its surface and 6.63% larger hemisphere when measuring its grey matter. Therefore, measures of grey matter volume may be more likely to capture interindividual differences of the *PT* asymmetry, whereas the surface measures may only scratch the top of the *PT* iceberg.

Interestingly, in a previous study in chimpanzees, Hopkins and Nir (2010) noted that leftward *PT* grey matter volume asymmetry constituted a better marker for the chimpanzee’s right-hand preference in communicative pointing gestures than *PT* surface (but see Meguerditchian et al., 2012). This latter study suggested the hypothesis that asymmetry of *PT* grey matter volume might be associated to functional asymmetry related to properties of gestural communication in apes, which have been found to share common features with human language such as intentionality, flexibility or referential properties (i.e., Liebal et al., 2013). Communicative manual gestures in baboons were also described in the literature (e.g., Molesti et al., 2020) as well as their chimpanzees-like manual lateralization patterns (Meguerditchian et al. 2013). Follow-up behavioral observations on gestural lateralization for communication in our sample of growing baboons will advance this question, once the focal subjects develop their full gestural repertoire. Specifically, taking advantage of the stronger *PT* asymmetries described in the present study for grey matter volume in comparison to surface measures, we could further investigate whether those early brain asymmetries might predict the gestural lateralization’s emergence in later development.

In conclusion, the present finding in nonhuman infants provides additional support to the hypothesis of a continuity between nonhuman and human primates concerning early leftward structural *PT* asymmetry in brain development. Early *PT* asymmetry might be thus not a human-specific marker for the pre-wired language-ready brain in infants. Nevertheless, it might be not excluded that this common anatomical signature is related to an ancient shared cognitive process at the heart of language evolution.

## Acknowledgments

We are very grateful to the vets Romain Lacoste and Marie Dumasy as well as the anesthetist Laura Laura Giacomino for supervising the first health and anesthesia monitoring, Emilie Rapha for great assistance and animal care, Frederic Charlin, as well as the care staff of the Station de Primatologie, such as Valérie Moulin, Brigitte Rimbaud, Richard Francioly, the vets Pascaline Boitelle, Alexia Cermolacce & Janneke Verschoor, the behavioral manager Pau Molina, the engineers of the MRI center, Bruno Nazarian and Jean-Luc Anton for coordinating the MRI sessions.

## Availability of data and material

See supplementary material

## Code availability

Not applicable

## Authors’ contributions

Y.B and A.M prepared the paper and the revision. R.P. performed the tracing and analyses. J.S. parametrized the MRI sequences and optimized the MRI acquisition setup. L.V. and L.R. designed and performed respectively the specific procedures of welfare, anesthesia, monitoring and preparation of baboons in the MRI machine. A.M. designed and supervised the study and MRI acquisitions.

## Ethics approval

All animal procedures were approved by the “C2EA-71 Ethical Committee of neurosciences” (INT Marseille) under the number APAFIS#13553-201802151547729 v4 and has been conducted at the Station de Primatologie under the number agreement C130877 for conducting experiments on vertebrate animals (Rousset-Sur-Arc, France). All methods were performed in accordance with the relevant French law, CNRS guidelines and the European Union regulations (Directive 2010/63/EU).

## References

Becker Y, Sein J, Velly L, et al (2021) Early Left-Planum Temporale Asymmetry in Newborn Monkeys (Papio anubis): A longitudinal structural MRI study at two stages of development. NeuroImage 117575. https://doi.org/10.1016/j.neuroimage.2020.117575

Borovsky A, Saygin AP, Bates E, Dronkers N (2007) Lesion correlates of conversational speech production deficits. Neuropsychologia 45:2525–2533. https://doi.org/10.1016/j.neuropsychologia.2007.03.023

Chi JG, Dooling EC, Gilles FH (1977) Left-Right Asymmetries of the Temporal Speech Areas of the Human Fetus. Archives of Neurology 34:346–348. https://doi.org/10.1001/archneur.1977.00500180040008

Dehaene-Lambertz G, Dehaene S, Hertz-Pannier L (2002) Functional Neuroimaging of Speech Perception in Infants. Science 298:2013–2015. https://doi.org/10.1126/science.1077066

Dronkers NF, Wilkins DP, Van Valin RD, et al (2004) Lesion analysis of the brain areas involved in language comprehension. Cognition 92:145–177. https://doi.org/10.1016/j.cognition.2003.11.002

Dubois J, Benders M, Lazeyras F, et al (2010) Structural asymmetries of perisylvian regions in the preterm newborn. NeuroImage 52:32–42. https://doi.org/10.1016/j.neuroimage.2010.03.054

Foundas AL, Bollich AM, Feldman J, et al (2004) Aberrant auditory processing and atypical planum temporale in developmental stuttering. Neurology 63:1640–1646. https://doi.org/10.1212/01.WNL.0000142993.33158.2A

Gannon PJ, Holloway RL, Broadfield DC, Braun AR (1998) Asymmetry of Chimpanzee Planum Temporale: Humanlike Pattern of Wernicke’s Brain Language Area Homolog. Science 279:220–222. https://doi.org/10.1126/science.279.5348.220

Geschwind N, Levitsky W (1968) Human Brain: Left-Right Asymmetries in Temporal Speech Region. Science 161:186–187. https://doi.org/10.1126/science.161.3837.186

Glasel H, Leroy F, Dubois J, et al (2011) A robust cerebral asymmetry in the infant brain: The rightward superior temporal sulcus. NeuroImage 58:716–723. https://doi.org/10.1016/j.neuroimage.2011.06.016

Greve DN, Van der Haegen L, Cai Q, et al (2013) A surface-based analysis of language lateralization and cortical asymmetry. J Cogn Neurosci 25:1477–1492. https://doi.org/10.1162/jocn_a_00405

Hill J, Inder T, Neil J, et al (2010) Similar patterns of cortical expansion during human development and evolution. Proceedings of the National Academy of Sciences 107:13135–13140. https://doi.org/10.1073/pnas.1001229107

Hopkins WD, Marino L, Rilling JK, MacGregor LA (1998) Planum temporale asymmetries in great apes as revealed by magnetic resonance imaging (MRI). NeuroReport 9:2913–2918

Hopkins WD, Nir TM (2010) Planum temporale surface area and grey matter asymmetries in chimpanzees (Pan troglodytes): The effect of handedness and comparison with findings in humans. Behavioural Brain Research 208:436–443. https://doi.org/10.1016/j.bbr.2009.12.012

Hopkins, W. D., Taglialatela, J. P., Meguerditchian, A., Nir, T., Schenker, N. M., & Sherwood, C. C. (2008). Gray matter asymmetries in chimpanzees as revealed by voxel-based morphology. Neuroimage, 42, 491–497.

Josse G, Hervé P-Y, Crivello F, et al (2006) Hemispheric specialization for language: Brain volume matters. Brain Research 1068:184–193. https://doi.org/10.1016/j.brainres.2005.11.037

Keller SS, Roberts N, García-Fiñana M, et al (2010) Can the Language-dominant Hemisphere Be Predicted by Brain Anatomy? Journal of Cognitive Neuroscience 23:2013–2029. https://doi.org/10.1162/jocn.2010.21563

Liebal, K., Waller, B. M., Burrows, A. M., & Slocombe, K. E. (2013). Primate communication: A multimodal approach. Cambridge: Cambridge University Press.

Lyn, H., Pierre, P., Bennett, A. J., Fears, S., Woods, R., & Hopkins, W. D. (2011). Planum temporale grey matter asymmetries in chimpanzees (Pan troglodytes), vervet (Chlorocebus aethiops sabaeus), rhesus (Macaca mulatta) and bonnet (Macaca radiata) monkeys. Neuropsychologia, 49, 2004–2012.

Manjón JV, Coupé P, Martí◻Bonmatí L, et al (2010) Adaptive non-local means denoising of MR images with spatially varying noise levels. Journal of Magnetic Resonance Imaging 31:192–203. https://doi.org/10.1002/jmri.22003

Marie D, Roth M, Lacoste R, et al (2018) Left Brain Asymmetry of the Planum Temporale in a Nonhominid Primate: Redefining the Origin of Brain Specialization for Language. Cereb Cortex 28:1808–1815. https://doi.org/10.1093/cercor/bhx096

Meguerditchian A, Gardner MJ, Schapiro SJ, Hopkins WD (2012) The sound of one-hand clapping: handedness and perisylvian neural correlates of a communicative gesture in chimpanzees. Proc Biol Sci 279:1959–1966. https://doi.org/10.1098/rspb.2011.2485

Meguerditchian A, Vauclair J, Hopkins WD (2013) On the origins of human handedness and language: A comparative review of hand preferences for bimanual coordinated actions and gestural communication in nonhuman primates. Developmental Psychobiology 55:637–650. https://doi.org/10.1002/dev.21150

Molesti, S., Meguerditchian, A., & Bourjade, M. (2020). Gestural communication in olive baboons (Papio anubis): repertoire and intentionality. Animal Cognition, 23, 19–40.

Ocklenburg S, Friedrich P, Fraenz C, et al (2018) Neurite architecture of the planum temporale predicts neurophysiological processing of auditory speech. Science Advances 4:eaar6830. https://doi.org/10.1126/sciadv.aar6830

Shapleske J, Rossell SL, Woodruff PWR, David AS (1999) The planum temporale: a systematic, quantitative review of its structural, functional and clinical significance. Brain Research Reviews 29:26–49. https://doi.org/10.1016/S0165-0173(98)00047-2

Spocter MA, Sherwood CC, Schapiro SJ, Hopkins WD (2020) Reproducibility of leftward planum temporale asymmetries in two genetically isolated populations of chimpanzees (*Pan troglodytes*). Proc R Soc B 287:20201320. https://doi.org/10.1098/rspb.2020.1320

Tzourio-Mazoyer N, Crivello F, Mazoyer B (2018) Is the planum temporale surface area a marker of hemispheric or regional language lateralization? Brain Struct Funct 223:1217–1228. https://doi.org/10.1007/s00429-017-1551-7

Vigneau M, Beaucousin V, Hervé PY, et al (2006) Meta-analyzing left hemisphere language areas: Phonology, semantics, and sentence processing. NeuroImage 30:1414–1432. https://doi.org/10.1016/j.neuroimage.2005.11.002

Wada JA (1975) Cerebral Hemispheric Asymmetry in Humans: Cortical Speech Zones in 100 Adult and 100 Infant Brains. Archives of Neurology 32:239. https://doi.org/10.1001/archneur.1975.00490460055007

Wernicke C (1874) Der aphasische Symptomencomplex: Eine psychologische Studie auf anatomischer Basis. Cohn.

Witelson SF, Pallie W (1973) Left Hemisphere Specialization for Language in the Newborn: neuroanatomical Evidence of Asymmetry. Brain 96:641–646. https://doi.org/10.1093/brain/96.3.641

Xia, J., Wang, F., Wu, Z., Wang, L., Zhang, C., Shen, D., Li, G., 2019. Mapping hemispheric asymmetries of the macaque cerebral cortex during early brain development. Hum. Brain Map. doi: 10.1002/hbm.24789.

