## Supplementary Material for "Planum Temporale grey matter volume asymmetries in new-born monkeys (*Papio anubis*)"

1) Supplementary figure of manual delineation procedure
2) Supplementary Table 1 (1^st^ longitudinal scan t0, subject details, N = 34)

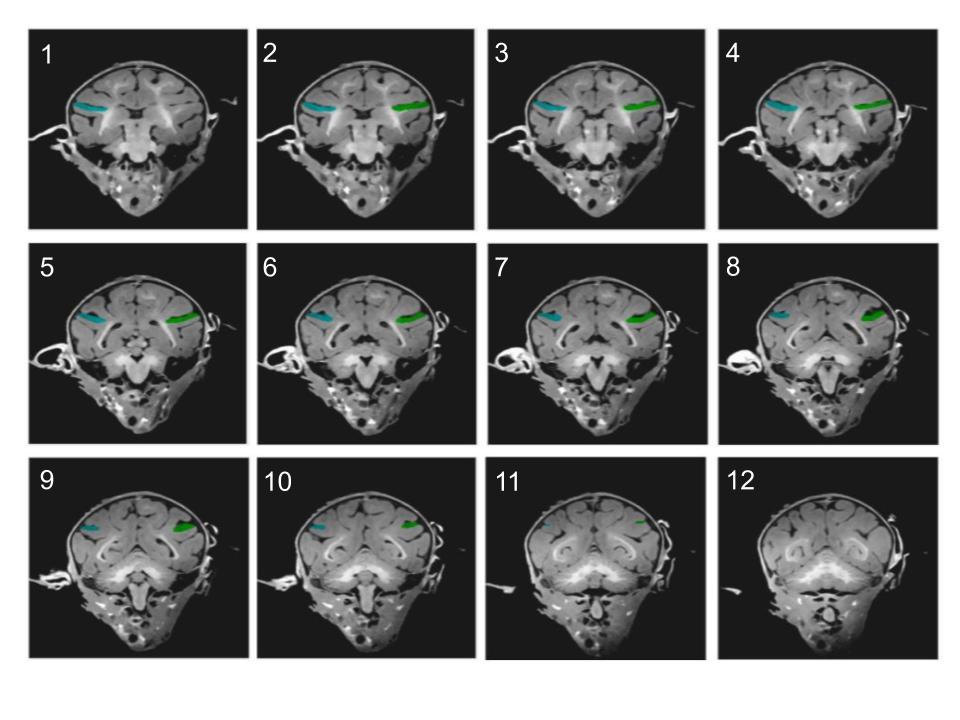
1) Supplementary figure of manual delineation procedure in both hemispheres of one newborn baboon brain, using coronal slices**.**
Shown below, 12 slices used for this tracing, slice 1 being the most anterior section used and slice 12, the most posterior. The extent of the left Planum Temporale volume is marked with a blue line in the left hemisphere and a green line in the right hemisphere.

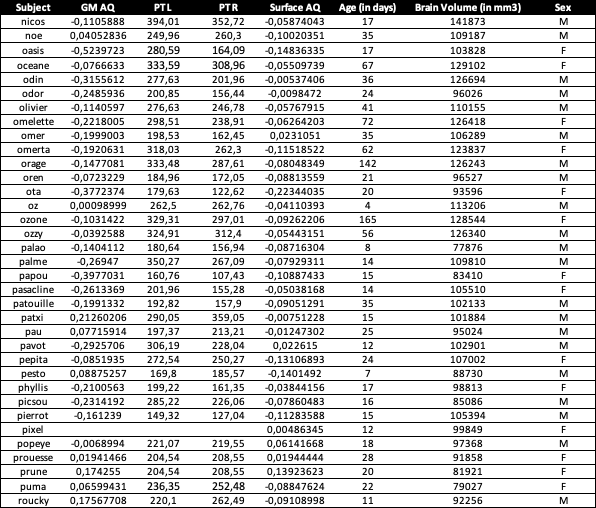
2) Supplementary Table 1 (1^st^ longitudinal scan t0, subject details, N = 34):
